# AlphaSimR: An R-package for Breeding Program Simulations

**DOI:** 10.1101/2020.08.10.245167

**Authors:** R. Chris Gaynor, Gregor Gorjanc, John M. Hickey

## Abstract

This paper introduces AlphaSimR, an R package for stochastic simulations of plant and animal breeding programs. AlphaSimR is a highly flexible software package able to simulate a wide range of plant and animal breeding programs for diploid and autopolyploid species. AlphaSimR is ideal for testing the overall strategy and detailed design of breeding programs. AlphaSimR utilizes a scripting approach to building simulations that is particularly well suited for modeling highly complex breeding programs, such as commercial breeding programs. The primary benefit of this scripting approach is that it frees users from preset breeding program designs and allows them to model nearly any breeding program design. This paper lists the main features of AlphaSimR and provides a brief example simulation to show how to use the software.

## Introduction

This paper introduces AlphaSimR, an R package for stochastic simulations of plant and animal breeding programs. Stochastic simulation is a powerful tool for design and optimization of breeding programs, because it provides a fast, inexpensive method for testing alternative breeding program designs. Simulations have been used to improve both plant breeding programs (e.g.; Lin et al. 2016; Gaynor et al. 2017; Gorjanc et al. 2018) and animal breeding programs (e.g.; Hayes and Goddard 2003; Jenko et al. 2015; Johnsson et al. 2019) as well as to address theoretical concepts in quantitative genetics and breeding (e.g., Gorjanc *et al*. 2015). AlphaSimR has been specifically designed to make simulations more common by providing an easy-to-use and highly flexible software package able to simulate a wide range of plant and animal breeding programs.

Stochastic simulations have rarely, if ever, been used to improve breeding programs for many agriculturally important species. This is likely due to the difficultly in setting up and running these simulations. This difficulty is in no small part due to the need for a person with thorough knowledge of breeding programs and computer programming. This person must possess a thorough understanding of the breeding programs they wish to simulate so that they can construct an informative simulation. They must also possess the programming skills needed to develop, run, and evaluate the simulations. The amount of programming skills this person needs to possess is considerable when there are not existing software programs for modeling the specific breeding program of interest. To address this issue, new software is needed that can lower the programming burden and thereby increase the ease of running simulations.

AlphaSimR has been specifically designed to make running stochastic simulations of whole breeding programs easier. To accomplish this goal, AlphaSimR provides the ability to run simulations both interactively or via scripts within the R software environment (R Core Team 2019). More specifically, AlphaSimR provides users with a range of R functions that correspond to common operations in a breeding program, such as crossing and selection. This allows users to apply functions representing breeding operations directly to objects that represent populations of animals or plants. The benefit of this approach is that it makes writing simulation code more intuitive, by allowing users to directly translate a description of a breeding program into an R script. It also provides a natural path for learning how to use the software by allowing users to start with simulations of simple breeding programs and gradually progress to more complicated breeding programs. With the respect to learning, simulations are also an invaluable tool to teach students and new professionals about theoretical and practical breeding concepts.

AlphaSimR is suitable for simulating a wide range of breeding programs and species. The software models the genomes of both diploid and autopolyploid species. The scripting approach employed by AlphaSimR allows for modeling nearly any breeding program structure, without limiting users to preset designs. AlphaSimR has been heavily optimized for large scale simulations (>1,000,000 individuals), because it is specifically designed for whole breeding program simulations.

## Methods

AlphaSimR is a large package with an extensive list of features, so we will only describe its main features here. For the sake of brevity, these descriptions are designed to provide an overview of AlphaSimR’s functionality and not a detailed accounting of its implementation. First, AlphaSimR’s approach to stochastic simulations will be given to provide a high-level overview of how the software works. Then, we will describe a few key elements of this approach before concluding with an overview of AlphaSimR’s implementation.

### Simulation approach

AlphaSimR uses a simulation approach that combines the coalescent and gene drop methods (Hickey and Gorjanc 2012). The coalescent method is used for backwards-in-time simulations. It is used in AlphaSimR to generate whole-chromosome founder haplotypes. The gene drop method is used for forwards-in-time simulations. It is used in AlphaSimR to create new haplotypes from the original founder haplotypes.

### Founder haplotypes

The preferred method for creating founder haplotypes in AlphaSimR is to use the Markovian Coalescent Simulator (MaCS; Chen *et al*. 2009). MaCS is included in AlphaSimR and used to generate founder haplotypes according to either a predefined parameter set for some species, or user supplied parameters. Alternatively, users can create founder haplotypes by importing external data into AlphaSimR or using built-in functions for random sampling. The option to import external data allows users to use other coalescent simulators or real genotypic data, provided the linkage phase and genetic map are known.

### Genetic recombination

AlphaSimR creates new haplotypes by modeling genetic recombination during meiosis. A genetic map is used to model the distribution of genetic recombination. AlphaSimR allows for sex-specific genetic maps to represent different recombination rates between sexes. The specifics for modeling meiosis in AlphaSimR depend on whether the species is a diploid or an autopolyploid.

For diploid species, AlphaSimR simulates meiosis and genetic recombination according to the gamma model (McPeek and Speed 1995). The gamma model accommodates crossover interference and has been shown to fit real data (e.g. Broman and Weber 2000). The magnitude of crossover interference is controlled by a single parameter that can be adjusted by the user.

For autopolyploid species, AlphaSimR simulates meiosis using a combination of bivalent and quadrivalent chromosome pairing. Bivalent or quadrivalent homologous pairs are chosen at random according to a parameter for the probability of quadrivalent pairing, which can be tuned by the user. Bivalent pairs are resolved using the gamma model for diploids. Quadrivalent pairs are resolved according to the model for “cross-type” configurations used in the PedigreeSim software (Voorrips and Maliepaard 2012). This model involves sampling chiasmata positions from a gamma distribution and resolving crossovers by sampling centromeres and working outwards towards the telomeres. This technique models unique features of meiosis in autopolyploids, such as recombinant chromosomes composed of three parental chromosomes and double reductions (Bourke *et al*. 2015).

### Traits

Traits in AlphaSimR are classified according to the biological effects they model. The biological effects modeled in AlphaSimR are: **A**dditive, **D**ominance, **E**pistatic, and **G**enotype-by-environment. The first letter of each effect is used to derive a name for each trait type under the **ADEG** framework. For example, a trait with only additive effects is called an **A** trait. A trait with additive and dominance effects is called an **AD** trait. AlphaSimR currently supports the following trait types: **A**, **AD**, **AE**, **AG**, **ADE**, **ADG**, **AEG**, and **ADEG**.

The modeling of biological effects is based on classic quantitative genetics models. For example, the additive effects are equivalent to additive effects described in a quantitative genetics textbook (e.g. Falconer and Mackay 1996). The modeling of the dominance effects allows for both directional dominance and a variable degree of dominance, ranging from partial dominance to overdominance (Gaynor *et al*. 2018). For autopolyploid species, the modeling of dominance represents digenic dominance. Epistatic effects are modeled as additive-by-additive epistatic effects between discrete pairs of loci. Genotype-by-environment effects are modeled as additive effects whose value is a function of a single environmental covariate.

AlphaSimR can simulate multiple traits using any combination of trait types. Each trait is simulated according to a user-defined number of QTL, which can differ between traits. Correlated traits can be simulated, provided they are pleiotropic and belong to the same trait type.

AlphaSimR uses a method for sampling QTL effects that is, to the authors’ knowledge, unique among stochastic simulation software. Users of AlphaSimR are asked to specify a desired mean and variance, either total or additive, for each trait. The software then samples QTL effects and scales the values for those effects to achieve precisely this mean and variance in a founder population. The benefit of AlphaSimR’s approach is that it allows users to set variables relating to the relative levels of dominance or epistasis independently of the founder population’s genetic variance. For example, a user can specify the average degree of dominance for QTL controlling a trait independently of the additive genetic variance for this trait.

### Variance components

AlphaSimR reports additive, dominance and additive-by-additive epistatic variances for any population. This is done without assuming random mating or linkage equilibrium, so that the values are correct regardless of the population’s genetic structure. This allows users to compare simulated populations to real-world data for the sake of benchmarking simulations. AlphaSimR also offers further partitioning of genetic variance into genic variance, covariance due to departures from Hardy-Weinberg equilibrium and covariance due to linkage disequilibrium, as described by Bulmer (1976).

### Selection

A wide range of functions are available for modeling selections. These functions allow for selection on multitude of criteria, such as: phenotypes, genetic values, breeding values, or estimated breeding values. Selection can be on one trait or an index of multiple traits. Selections can also be modeled as selection between or within families or over an entire population. AlphaSimR also supports user supplied selections, allowing users to implement their own selection methods, for example optimum contribution selection as in Gorjanc et al. (2018)

### Mating and propagation schemes

A wide range of functions are available in AlphaSimR for modeling common mating and propagation schemes. These schemes include: biparental crossing, selfing, clonal propagation, generation of doubled haploid lines, and propagation in open pollinating populations with variable degrees of selfing. AlphaSimR also supports user supplied mating plans.

### Genomic prediction

Modeling genomic prediction in breeding programs is one of the main use cases for AlphaSimR. AlphaSimR offers several built-in functions for fitting common genomic prediction models. The built-in functions use mixed model solvers based on the following R packages: rrBLUP (Endelman 2011), EMMREML (Akdemir and Godfrey 2015) and Sommer (Covarrubias-Pazaran 2016). Each solver has been optimized for performance within AlphaSimR and written in C++ using the R packages Rcpp (Eddelbuettel and Francois 2011) and RcppArmadillo (Eddelbuettel and Sanderson 2014). Users can also make use of other R packages or external applications for modeling genomic prediction. This is done by exporting data from an AlphaSimR simulation into another R function or external program for genomic predictions, generating predictions, and importing the predictions back into AlphaSimR objects.

### Implementation

Much of AlphaSimR’s code has been written in C++ to improve performance. For example, this has been used to implement bitwise storage of genotype data to reduce memory usage and enable multithreading for increased speed. AlphaSimR also improves performance by limiting data storage and calculations, such as variance component calculations, to only those expressly requested by the user. This approach differs from other stochastic simulation programs, including the original AlphaDrop (Hickey and Gorjanc 2012) and AlphaSim (Faux *et al*. 2016), which typically perform all calculations and save all data.

## Results and Discussion

### Example Simulation

This section will demonstrate AlphaSimR using a simulation of a single breeding cycle for a generic wheat breeding program. The code needed to run this simulation is presented below after a brief description of the breeding program.

Figure 1 shows a schematic representing the stages of the generic wheat breeding program with a seven-year breeding cycle. In the first year, 200 bi-parental populations are produced by crossing and production of doubled haploid (DH) lines from those bi-parental populations begins. In the second year, the production of DH lines is completed in. In the third year, the DH lines are visually evaluated in a head-row (HDRW) nursery. In the remaining years, lines are selected based on performance in the previous year and evaluated in a yield trial. The yield trials are conducted over the course of three years before selecting a variety to release.

**Figure 1.**
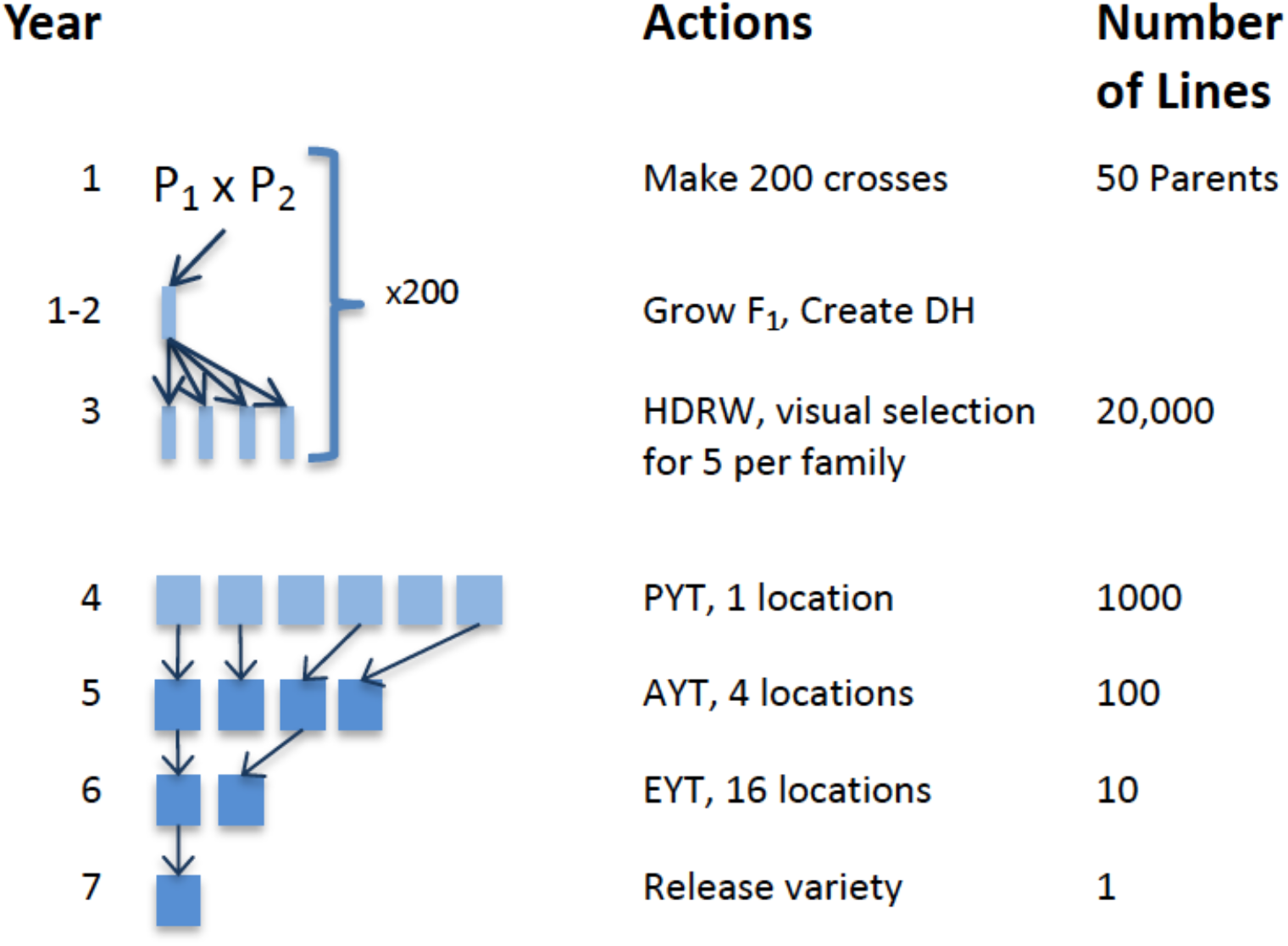
An overview of the variety development cycle for the example wheat breeding program. A variety is developed over the course of seven years. The steps in the development cycle are: making bi-parent crosses, forming doubled haploid (DH) lines, visually select lines grown in headrows (HDRW), evaluate lines in a preliminary yield trial (PYT), evaluate lines in an advanced yield trial (AYT), evaluate lines in an elite yield trial (EYT), and release a variety.

The first step is to generate founder haplotypes using MaCS. The founder haplotypes will be used to form the initial parents in the breeding program. Code for simulating the founder haplotypes for 50 inbred individuals is shown below. Each individual will have 21 chromosomes, each with 1000 segregating sites.

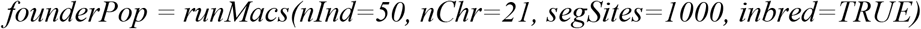

The second step is to set global parameters. Below is code for setting simulation parameters to model a single trait. The trait models additive genetic effects on 1000 loci per chromosome. The trait is also modeled as having a broad-sense heritability of 0.4 for evaluation in a single location.

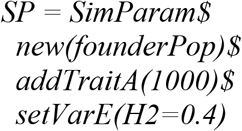

The next step is to simulate each year of the breeding program. In the first year, 200 bi-parental populations are produced by crossing the parents formed from the founder haplotypes. This code is presented below. The first line uses the founder haplotypes to form the parents and the second line makes 200 randomly chosen crosses between those parents.

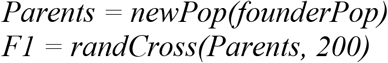

In the second and third years, the DH lines are produced and then they are evaluated in the HDRW nursery. The code for both these years is presented below. The first line forms 100 DH lines per F_1_ plant. The second line models evaluation in the HDRW nursery for the previously defined additive trait. The broad-sense heritability of this trait is reduced to 0.1 to represent visual selection.

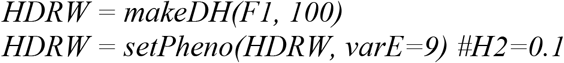

In the fourth year, the best entries in the HDRW nursery are selected and evaluated in a preliminary yield trial (PYT). This is modeled with the code below. The first line models selection in the HDRW by selecting the best lines within families. The second line models evaluation of the PYT at one location. The accuracy of this evaluation is based on the broad-sense heritability defined in the simulation parameters.

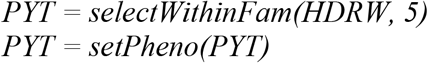

In the fifth year, the best PYT entries are selected and evaluated in an advanced yield trial (AYT). This is modeled with the code below. The first line models selection of the best PYT lines. The second line models evaluation of the AYT at four locations, which are represented as reps in the code.

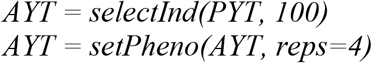

In the sixth year, the best AYT entries are selected and evaluated in an elite yield trial (EYT). This is modeled with the code below. The first line models selection of the best AYT lines. The second line models evaluation of the EYT at sixteen locations.

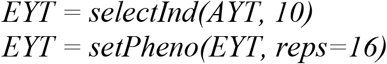

In the seventh year, the best performing EYT entry is chosen for release as a variety. This is modeled with the code below.

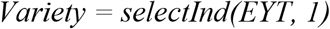

The final step is to evaluate the simulation results. This is done by producing a boxplot for the genetic values of entries in stage of the breeding program. The boxplot is shown in Figure 2. The code for generating the boxplot is given below. The first line of code extracts the genetic values for each entry and saves it in a list. The second line creates the boxplot showing the distribution of genetic values for entries in each stage of the breeding program.

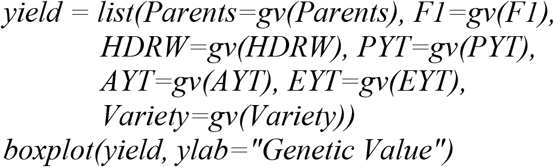

**Figure 2.**
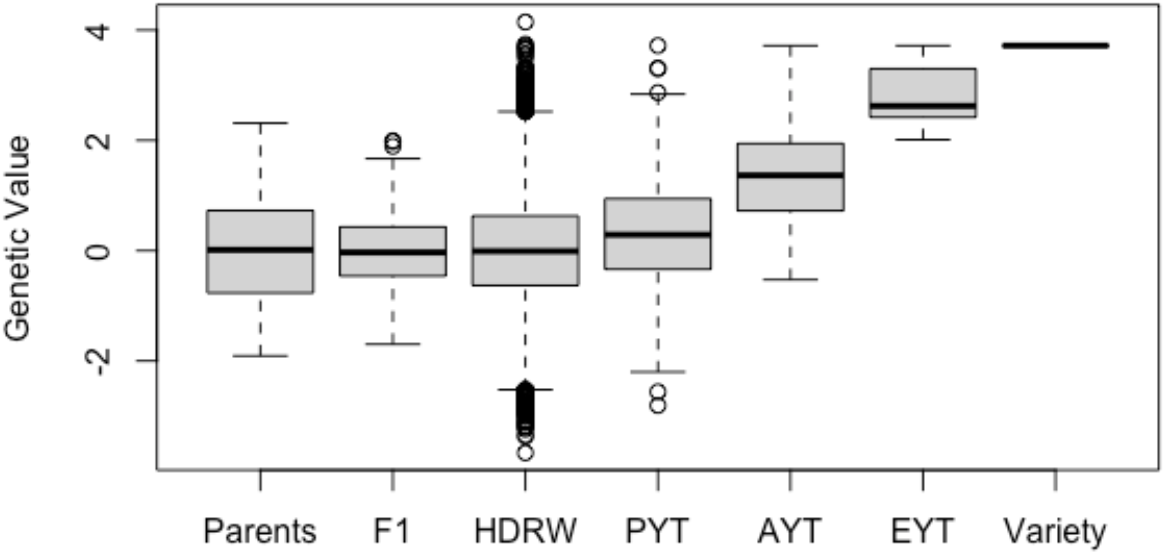
The distribution of genetic values in one replicate of the example breeding program. Separate boxplots are given for each stage of the breeding program.

### Concluding remarks

AlphaSimR represents a considerable improvement over its predecessor in terms of ease-of-use, flexibility, and computational efficiency (AlphaSim; Faux *et al*. 2016). It has been used in a handful of published simulations (Gorjanc *et al*. 2018; Muleta *et al*. 2019; Johnsson *et al*. 2019) as well as numerous unpublished simulations. The largest simulation undertaken in AlphaSimR to date involved over a hundred million individuals (unpublished), a feat that would not be feasible with original AlphaSim.

The improvements made to AlphaSimR make it uniquely well suited for simulating whole breeding programs. These types of simulations serve as a valuable tool for aiding strategic decision making within breeding programs. For example, AlphaSimR can be used test the economic value of modifying an existing breeding program. This will be of particular interest to breeding programs considering implementing genomic selection or changing their current implementation. These types of simulations can also be used to optimize selection stages or compare the efficiency of mating strategies.

AlphaSimR can be used for a wide range of simulations outside of whole breeding program simulations. For example, AlphaSimR can be used to test QTL mapping strategies or marker imputation strategies. AlphaSimR is also well suited for running simulations that help with teaching quantitative genetics and breeding. This is because students can be quickly taught how to use AlphaSimR for simple simulations, and the software’s ability to report variance components, perform genomic evaluations and evaluate accuracy of evaluations against the simulated true values is highly instructive.

AlphaSimR is under continuous development with new features being added on a semi-regular basis. Additional planned features include developing standard breeding program blueprints for major species and developing easy-to-use graphical user interfaces for these blueprints. These planned additions should make AlphaSimR even more user-friendly than it currently is.

### Web resources

AlphaSimR is publicly available on CRAN (https://CRAN.R-project.org/package=AlphaSimR). Additional documentation as well as links to graphical user interfaces for specialized applications are available on the AlphaGenes website (https://alphagenes.roslin.ed.ac.uk/wp/software-2/alphasimr/). A repository of example simulation scripts for learning to use the software, modeling specific breeding programs, and learning quantitative genetics principles are available on Bitbucket (https://bitbucket.org/hickeyjohnteam/alphasimr_examples).

## Acknowledgments

The authors acknowledge the financial support from Bayer Crop Science and the Gates Foundation.

